# Bovine serum albumin as an immunogenic carrier facilitating the development of hapten-specific monoclonal antibodies

**DOI:** 10.1101/2020.11.25.397455

**Authors:** Ruotong Zhao, Mingjun Jiang

## Abstract

There is a common misconception that the generation of hapten-specific monoclonal antibodies (mAbs) requires the use of a heterologous conjugate to ensure carrier-specific antibodies not being detected. In this study, salbutamol (SAL) was used as a model hapten to exhibit the benefits of bovine serum albumin (BSA) as a carrier for developing hapten-specific mAbs. SAL-BSA conjugate would serve as both an immune antigen and a screening antigen during the preparation of SAL-specific mAbs. Six hybridomas were identified to secret mAbs specific for free SAL with minor or negligible cross-reactivity with other β-agonists. Meanwhile, none of hybrodomas secreting anti-BSA antibodies were screened out even though the fetal bovine serum (FBS) added to the medium decreased from 10% to 1% (v/v). Based on one of the six mAbs, 3F12, a direct competitive enzyme-linked immunosorbent assay (dcELISA) was developed for meausring SAL. Under the optimized assay, the quantitative working range was from 312.5 to 20,000 pg/mL (*R*^2^ = 0.9959), with a limit of detection (LOD) of 142.9 pg/mL. The results showed that BSA is an efficient and suitable protein carrier for facilitating the development of hapten-specific mAbs.

## 1. Introduction

To develop specific and sensitive immunoassays for small-molecule compounds (haptens) detection, the crucial step is to generate high quality antibodies against the target molecule (Chappey et al. 1992). Haptens, however, are usually nonimmunogenic unless they are conjugated with an immunogenic carrier. In the majority of hapten-carrier systems, the B cells will produce antibodies that are specific for both the hapten and carrier (Parker 2013). Accordingly, different carrier proteins required to be used in immunization and screening/purification for screening hapten-specific rather than carrier-specific antibodies (Tong et al. 2015; Zhang et al. 2007). Despite of numerous reviews outlining the theoretical advantages of monoclonal antibodies (mAbs) over polyclonal antibodies (pAbs), most competitive enzyme immunoassays for haptens were based on pAbs (Goodrow and Hammock 1998; Lei et al. 2010; Darwish et al. 2012) since production of mAbs requires time-consuming and labor-intensive multistep screening procedures with heterologous conjugate (Chappey et al. 1992; Zhang et al. 2009). Therefore, eliminating the preparation of heterologous conjugates and simplifying the screening procedure would promote the development of hapten-specific antibodies.

Keyhole limpet hemocyanin (KLH) (Pak et al. 1995; Ling et al. 2014), bovine serum albumin (BSA) (Kim et al. 2004; López-Moreno et al. 2014) and ovalbumin (OVA) (Green et al. 2012; Parra et al. 2014) are the three most common carrier proteins employed for hapten conjugation. In many cases, KLH (or BSA) conjugate is preferred as imunogen for immunization, while BSA (or OVA) conjugate is used as coating antigen to monitor the raised antibodies (Jin et al. 2014; Saeed et al. 2017; Gao et al. 2007; Esteve-Turrillas et al. 2015). Hence, BSA or OVA, is often used as control carrier to verify that the antibodies are specific for the target hapten rather than the carrier itself (e.g. KLH or BSA). In order to avoid undesired cross-reactivity with the linker regions in antibody-screening procedures and final applications, moreover, different coupling strategies must be employed for preparing immunogens and coating antigens (Esteve-Turrillas et al. 2010; Vasylieva et al. 2015).

In our opinion, BSA is the most suitable carrier protein for preparing mAbs against haptens. Thus, the present study was an attempt to use salbutamol (SAL), a β-adrenergic receptor agonist illegally used for feed additive with a molecular weight of 239.31, as a model hapten to show the advantages of BSA as a carrier protein for preparation of hapten-specific mAbs.

## 2. Materials and methods

### 2.1 Reagents

1-Ethyl-3-(3-dimethylaminopropy) carbodiimide (EDC), bovine serum albumin (BSA), ovalbumin (OVA) horseradish peroxidase (HRP), N-hydroxysuccinimide (NHS), Freund’s complete and incomplete adjuvants, polythylene glycol (PEG, Mw 1450), HT and HAT Media Supplement were supplied by Sigma-Aldrich (Madrid, Spain). Dulbecco’s modified Eagle’s medium (DMEM) and fetal bovine serum (FBS) were from HyClone Laboratories (Logan, UT, USA). Salbutamol sulfate (SAL), ractopamine hydrochloride, clenbuterol hydrochloride and phenylethanolamine A were from Green Stone Switzerland Co., Ltd. All other chemicals and organic solvents were of analytical grade or better.

### 2.2 Conjugates of SAL-BSA and SAL-OVA preparation

The introduction of carboxylic group into the structure of SAL, required for protein conjugation, was achieved as described by Beaulieu N, et al. (Beaulieu et al. 1985) with a slight modification. In brief, 5 mg of succinic anhydride was added dropwise to the solution of 10 mg SAL in 5 mL at room temperature (RT), and the reaction mixture became orange in color. After centrifugation, the precipitates were thoroughly washed three times with ethanol to remove the unreacted SAL and succinic anhydride, and dried under a nitrogen atmosphere to yield about 15 mg of yellow amorphous solid.

Conjugates of SAL-BSA and SAL-OVA were prepared according to the mixed anhydride method (Erlanger et al. 1957) In this procedure, 5 mg of the dry derivative was dissolved in 2 mL of distilled water followed by the dropwise addition of 1.4 mg of EDC and 1 mg of NHS with constant stirring. The reaction mixture stood at RT for 24 h. Subsequently, BSA or OVA (10 mg) dissolved in 5 mL of 0.13 M sodium bicarbonate solution was dropwise added to the above solution with continuous stirring, and then further stirred at RT for 24 h. The conjugates were then purified by dialysis against PBS for one day with two changes of fresh buffer. Insoluble precipitates were removed by centrifugation (3,000 rpm, 10 min). The dialyzed solutions were frozen at –20 °C until use. Ultraviolet (UV) and sodium dodecyl sulfate polyacrylamide gel electropheresis (SDS-PAGE) were used to identify the conjugates.

### 2.3 Immunization and antibody generation

Seven-week-old BALB/c female mice (Comparative medicine center, Yangzhou University, Yangzhou, China) were vaccinated intraperitoneally with 0.2 mL of an emulsion containing Freund’s complete adjuvant and SAL-BSA conjugate (1:1; at a dose of 5 μg SAL-BSA conjugate per mouse). One booster vaccination was given in two weeks later except for the conjugate emulsified in Freund’s incomplete adjuvant.

Serum was collected ten days later and tested for antibody titres by indirect enzyme-linked immunosorbent assay (iELISA) and indirect competitive ELISA (icELISA) to determine specific IgM or IgG-class antibody for SAL-OVA and free SAL, respectively.

### 2.4 Cell fusion and screening

Three days before cell fusion, the mouse was challenged ip with SAL-BSA conjugate alone (0.1 mg/mL in PBS, 0.2 mL/mouse). Splenocytes from the immunized mouse were fused with SP2/0 myeloma cells according to standard protocols. Fused cells were resuspended gently in 100 mL of DMEM containing 15% FBS and 1× HAT media supplement, and evenly distributed into 96-well cell plates and incubated at 37°C in 5% CO_2_ (day 0). At day 5, half of the medium was replaced with fresh DMEM containing 10% FBS and 1× HT media supplement. At day 7, fresh DMEM containing 10% FBS and 1× HT media supplement completely substituted for the medium.

The antibody-producing hybridoma were screened by a two-step procedure. On the 10th day of fusion, hybridoma culture supernatants were screened by iELISA. The antibody secreting hybridoma cells were expanded into 48-well plates for being further confirmed to be SAL-specific by indirect competitive ELISA (icELISA). Hybridomas with desired characteristics were subcloned and expanded for antibody production by being injected intraperitoneally into the pristine primed BALB/c mice to obtain ascetic fluids.

### 2.5 First step: iELISA screening

The 384-well microplates were coated with 10 μL/well of SAL-BSA or SAL-OVA at concentration of 10 μg/mL in carbonate-buffered saline (CBS, 0.05 M, pH 9.6) at 37°C for 2 h. After washing three times with phosphate-buffered saline (0.01 M, pH 7.4) containing 0.05% Tween-20 (PBST), the culture supernatants (10 μL/well) were respectively added into the SAL-BSA conjugate coated plates and SAL-OVA conjugate coated plates, and incubated at 37°C for 1 h. The fresh complete medium was used as negative controls. After another washing, the plates were incubated at 37°C for another 1 h with HRP conjugated goat anti-mouse IgG and IgM antibodies diluted 1/10,000 in PBST (10 μL/well). The final washing procedure was followed by a color development, which was initiated by adding 10 μL/well of 3,3′,5,5′-Tetramethylbenzidine liquid substrate to measure the HRP tracer activity. After incubating at 37°C for 30 min, the enzymatic reaction was terminated by adding 10 μL/well of 1 M H_2_SO_4_, and the absorbance was measured at 450 nm with a Multiskan Spectrum (Thermo Electron Corporation, San Jose, CA, USA).

### 2.6 Second step: icELISA screening

Each well of 96-well microplates was coated with 100 μl of SAL-BSA conjugate (1.0 μg/mL) diluted with SCB at 37°C for 2 h. After washing three times with PBST, 50 μL of SAL (1.0 μg/mL diluted in PBS) and 50 μL of the supernatants of positive hybridomas were respectively added to each well. The subsequent steps were performed as above. Relative absorbance was calculated using the formula 100 × B/B0, where B and B0 were the absorbance of the well with and without SAL. The relative absorbances are around 100% for antibodies not reacting with free SAL.

### 2.7 Effect of FCS contents in culture medium on anti-BSA antibody detection

During the experiment, 96 SAL-BSA-negative hybridoma, each hybridoma divided into three equal parts, were transferred from 96-well cell plates to 48-well cell plates for further BSA-specific antibody screening. On the 3rd day, the medium was replaced with the medium containing 1%, 2% or 5% of FBS. After that the hybridoma cells would be allowed to grow until all died, and the culture supernatants were collected for BSA-specific antibody detection by iELISA as above with BSA as coating antigen.

### 2.8 Characterization of SAL-specific mAbs

The ascetic fluids were obtained and purified by the caprylic acid/ammonium sulfate method of McKinney and Parkinson (McKinney and Parkinson 1987). The protein content was determined by BCA kit (Pierce, USA). The mAbs were isotyped using a mouse mAb isotyping test kit (Sigma-Aldrich, USA). The icELISA described above was used to evaluate the affinity (Ka) and sensitivity of each mAb. The median inhibitory concentration (IC_50_) values were calculated by Graph pad Prism 5 to determine the sensitivity. Specificity was defined as the ability of structurally related analogs, including ractopamine, clenbuterol and phenylethanolamine A, to bind the specific mAb and the CR was calculated as: (IC_50_ of SAL)/(IC_50_ of analogs) × 100.

### 2.9 A direct competitive ELISA for SAL

After checkerboard optimization of the SAL-BSA coating amount and HRP-conjugated antibody concentration, a direct competitive ELISA (dcELISA) was developed for SAL using SAL-BSA conjugate at 10 μg/mL and HRP-conjugated mAb 3F12 at 1:2,500. Results were reported as the mean of two separate light absorbance values subtracted from the negative control value.

### 2.10 Statistical Analysis

Mean and standard deviation (SD) were calculated using Microsoft Excel 2007 software. All values in the text, tables, and figures were presented as mean ± standard deviation.

## 3. Results

### 3.1 Conjugates of SAL-BSA and SAL-OVA verification

UV spectra and SDS-PAGE were employed to obtain evidences of successful conjugation of SAL to proteins. Full-scan experiments over the range of 230-390 nm were performed to analyze the UV absorption characteristics of SAL, BSA, OVA and the conjugates. UV spectra in Fig. 1A and 1C showed a red-shift at 280 nm for SAL-BSA compared with 278 nm for BSA, and a red-shift at 279 nm for SAL-OVA compared with 277 nm for OVA. Analysis of SAL conjugates by SDS-PAGE (Fig. 1B and 1D) revealed an increase in the molecular weight of the conjugates when compared with BSA and OVA, respectively. These results clearly indicate the successful conjugation of SAL to BSA and OVA, respectively.

**Fig. 1.**
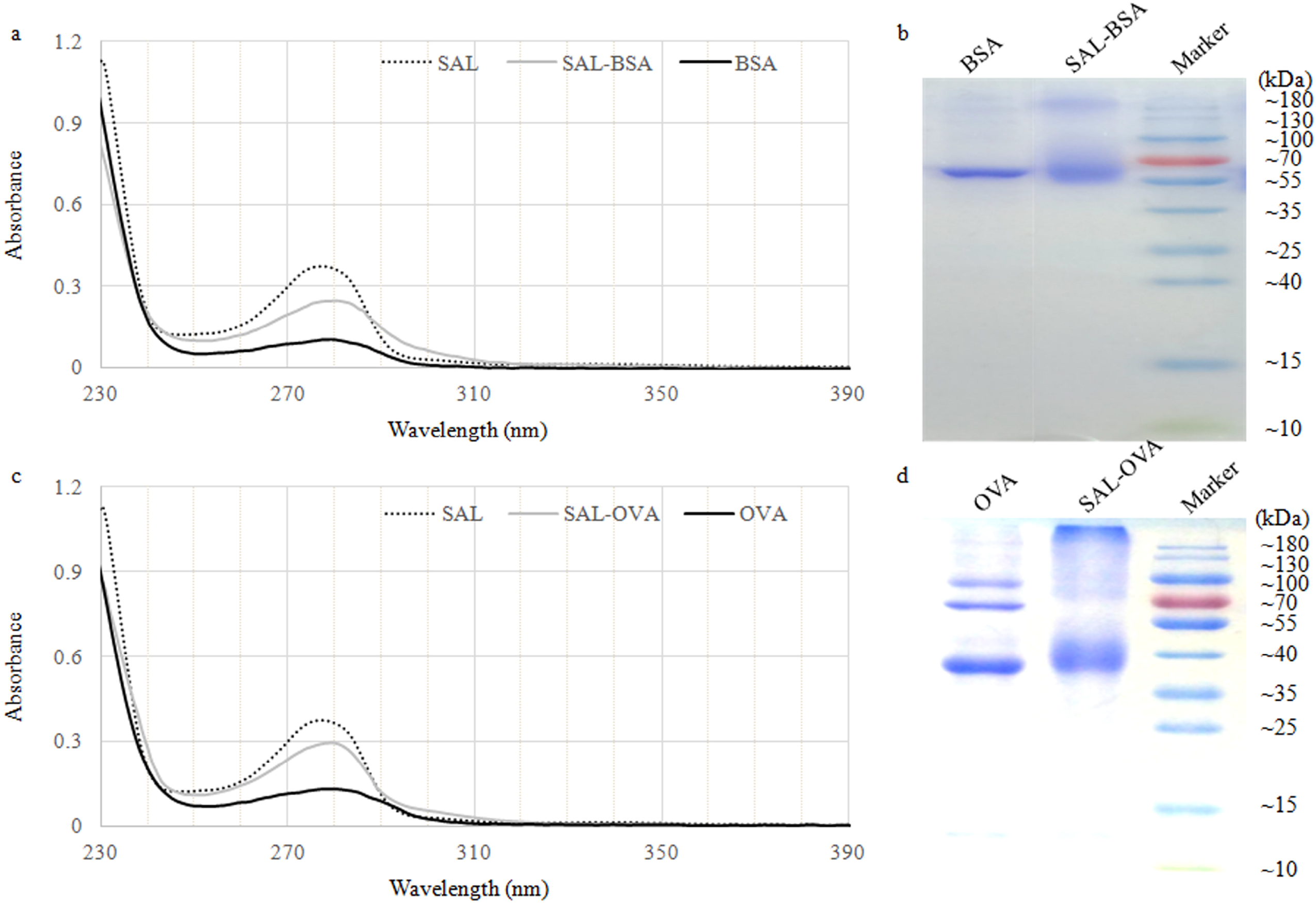
UV spectra and SDS-PAGE analysis of the conjugates of SAL-BSA and SAL-OVA Comparative UV absorption spectra with the conjugate and controls. Two microliters of the samples prepared in PBS at about 0.2 mg/ml were injected into NanoDrop 2000 spectrophotometer (Thermo Fisher Scientific Inc. USA) (a) and (c). The SDS-PAGE analysis of the conjugates stained with Coomassie brilliant blue. The SDS-PAGE of the conjugates and carrier proteins were performed using Mini-PROTEAN Tetra System (Bio-Rad Laboratories, Inc. USA) (b) and (d). The SDS-PAGE procedure was performed as previously described by Sambrook 15% separated gel and 5% concentrated gel was used (Sambrook and Russell 2006).

### 3.2 Determination of SAL-specific antibody titres

Anti-SAL IgM and IgG antibody responses were determined ten days after the third immunization and typical antibody dilution curves were shown in Fig. 2. The antiserum had an IgM antibody binding activity to the SAL-OVA conjugate of 2.049 at 1:1,000 and bound at a dilution of up to 1:32,000. However, after competition with free SAL, the activity was reduced to 1.330 at 1:1,000, and the serum reacted at a dilution of up to 1:32,000 (Fig. 2A). Similar results were obtained with IgG. The antiserum without SAL had an antibody binding activity of 2.642 at a dilution of 1:1,000 and reacted at dilutions of up to 1:32,000. After competition, the binding activity was reduced to 1.479 at a 1:1,000 dilution and bound at a dilution of up to 1:32,000 (Fig. 2B). These results show a successful immunogenic response induced by SAL-BSA conjugate.

**Fig. 2.**
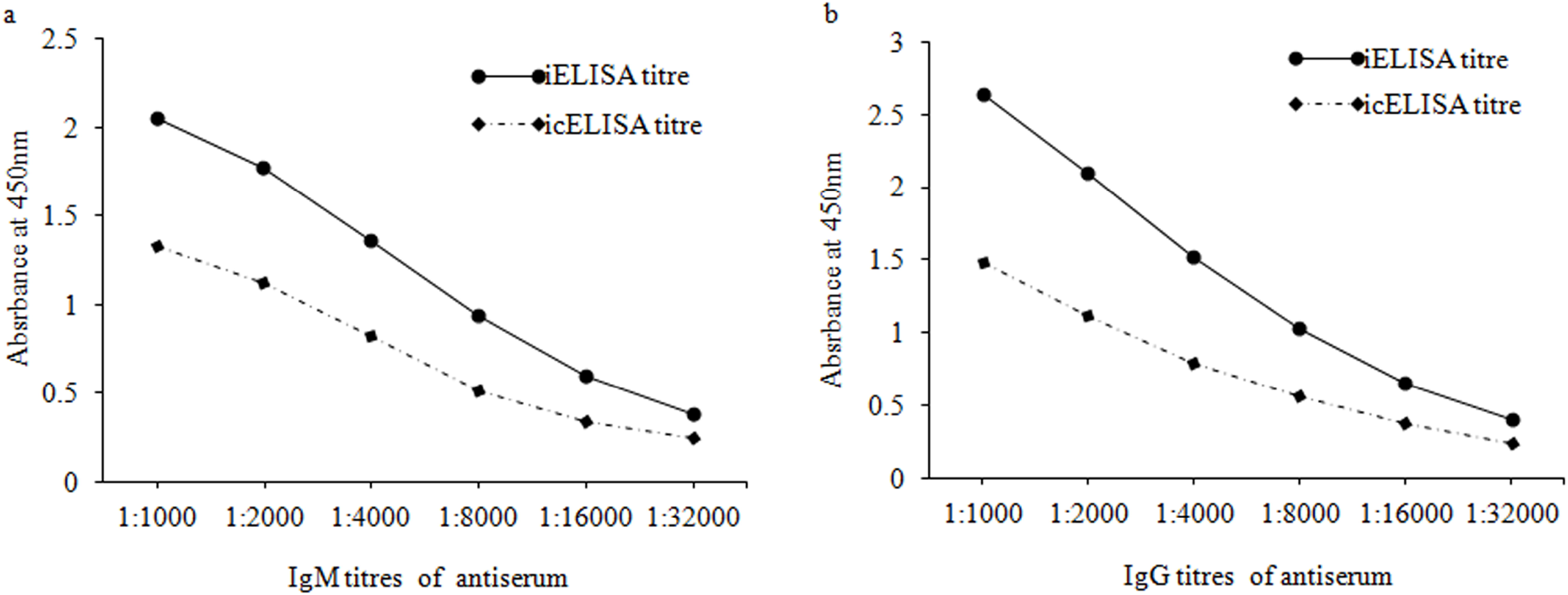
Dilution curve of a mouse antiserum raised against the SAL-BSA conjugate The specific titres of IgM (a) and IgG (b) to the SAL-BSA conjugate and SAL alone were assayed with serial dilutions of antiserum by iELISA and icELISA, respectively. The iELISA and icELISA were performed as described in “Materials and Methods” except the serial dilutions of antiserum instead of the supernatants of hybridomas.

### 3.3 SAL-specific hybridoma screening

Following fusion of splenocytes with SP2/0 cells and outgrowth of primary hybridoma, the hybridoma culture supernatants from 351 wells were evaluated by iELISA on the 10th day after fusion. Fifteen hybridomas culture supernatants were found to contain antibody-specific to SAL-BSA conjugate, which was consistent with that of SAL-OVA conjugate as coating antigen. After expansion into 48-well plates, nine primary hybridomas were still positive for anti-SAL-BSA antibodies confirmed by iELISA. The positive hybridomas were subsequently tested by icELISA using SAL as competitive inhibitor (1 μg/mL). Eventually, five hybridomas secreting mAbs specific for free SAL were cloned and stabilized (Table 1). The mAbs derived from the rest four hybridomas targeted the linker region of SAL-BSA due to their binding activity being only partially or negligibly inhibited by free SAL.

**Table 1.**
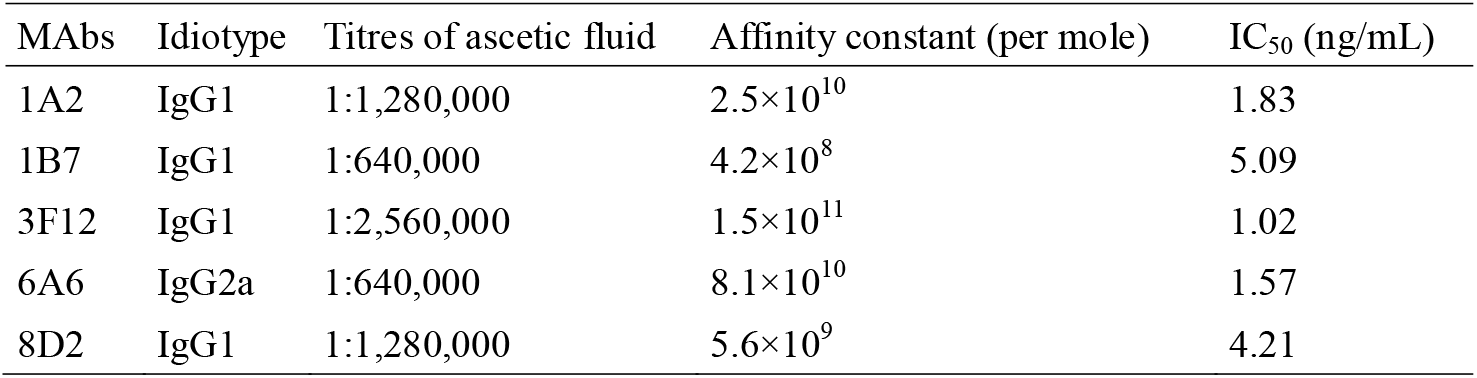
Characteristics of anti-SAL mAbs

### 3.4 BSA-specific hybridoma screening

A total of 96 hybridoma clones were expanded into 48-well plats with the medium decreased in FBS contents from 5% to 1% (v/v). No hybridoma was found to produce BSA-specific mAbs.

### 3.5 Characterization of SAL-specific mAbs

The characteristics of five mAbs in terms of titre, affinity, class, and subclass were summarized in Table 1. Except for mAb 6A6 belonging to IgG2a, all other mAbs were IgG1 with λ light chain, and the affinity constants of the mAbs ranged from 10^8^ to 10^11^ per mole. The IC50 values of the mAbs for SAL ranged from 1.02 to 5.09 ng/mL with mAb 3F12 being the lowest. Therefore, the specificity of mAb 3F12 was further evaluated using the other β-adrenergic receptor agonists, and showed that the CRs were 0.51%, 31.4% and 0.92% for ractopamine hydrochloride, clenbuterol hydrochloride and phenylethanolamine A, respectively. The mAb 3F12 was conjugated to horseradish peroxidase (HRP) through the periodate method (Tijssen and Kurstak 1984).

### 3.6 Determination of SAL by dcELISA

After checkerboard optimization of the coating antigen (SAL-BSA) and detection antibody concentration (HRP-3F12), a representative standard curve for the dcELISA was depicted in Fig. 3. A strong negative linear relationship between the relative absorbance and log concentration of SAL was obtained in the range of 312.5–20,000 pg/mL (*R*^2^ = 0.9959). The limit of detection (LOD) was 142.9 pg/mL of SAL (LOD was calculated as mean – 3× SD, where mean was based on the responses of blank samples and SD was the standard deviation of analytical responses.). The limit of quantitation (LOQ) was 193.9 pg/mL SAL (mean – 10× SD, n = 3).

**Fig. 3.**
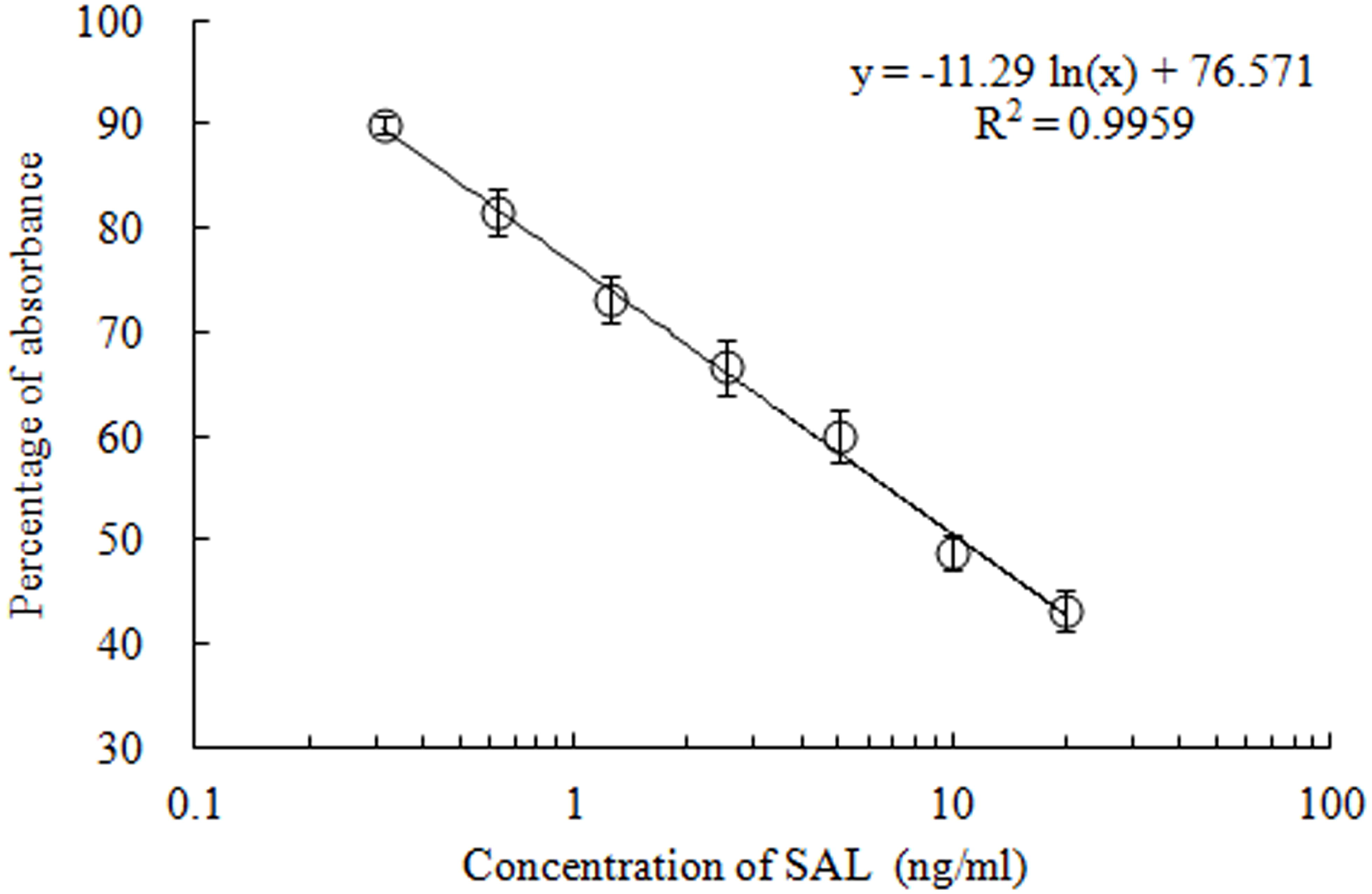
The standard curve for SAL in PBS was determined by the dcELISA Vertical bars represent standard deviations for three independent assays.

Intra- and inter-assay precisions were qualified with three standard solutions of SAL at 625, 2,500 and 10,000 pg/mL, which was calculated by the relative standard deviation (RSD) resulting from well to well (n = 6) within one plate, and inter-assay precision obtaining from different plates (n = 3). The analytical performance of the assay was demonstrated in Table 2. Precision analysis showed low RSD (≤ 9.7%) and good accuracy (95.6–108.8%).

**Table 2.** Intra-assay and inter-assay precisions of SAL detection by the dcELISA

| Assay | Standards (pg/mL) | Mean ± SD | RSD (%) | Accuracy (%) |
| --- | --- | --- | --- | --- |
| Intra-assay | 625 | 665.0 ± 35.1 | 5.3 | 106.4 |
| | 2,500 | 2,489.7 ± 110.9 | 4.5 | 99.6 |
| | 10,000 | 10,879.2 ± 243.5 | 2.2 | 108.8 |
| Inter-assay | 625 | 645.0 ± 62.5 | 9.7 | 103.2 |
| | 2,500 | 2,390.2 ± 163.1 | 6.8 | 95.6 |
| | 10,000 | 10,199.5 ± 535.8 | 5.3 | 102.0 |

## 4. Discussion

A hapten is a small molecule that can elicit an immune response only when covalently attached to a large carrier such as a protein. As an ideal carrier of hapten, single polypeptide protein BSA has numerous functional groups (amines, hydroxyls, carboxyls, or sulfhydryl groups) for effective conjugation. ^(12, 13)^ In addition, BSA has been proved to be heterologous enough to evoke an adequate immune response in mice (Hirayama et al. 1990).

During immunization with hapten-BSA conjugates, the immunized mice will produce antibodies against both the hapten and BSA. In order to avoid the false-positive results in screening tests, there is a misconception that heterologous conjugates are necessary for screening of hapten-specific antibody-secreting hybridoma. Accordingly, the hepten-OVA conjugates, for instance, should be used to detect the hapten-specific antibodies, and vice versa (Jin et al. 2014; Saeed et al. 2017; Gao et al. 2007; Esteve-Turrillas et al. 2015; Esteve-Turrillas et al. 2010). However, in the present study, no hybridoma secreting anti-BSA mAb was screened out using the SAL-BSA conjugate for antibody screening, even though the FBS content in the supernatant was reduced from 10% to 1.0%. Typically, the mAb content in the supernatant of hybridoma is less than 10 μg/mL, which is equivalent to 70 nmol mAb per liter (Kuhne et al. 2014; Tomita et al. 2011). Meanwhile, the average content of albumin in FBS is about 23g/L, which is equivalent to 350 μmol BSA per liter of the hybridoma culture supernatant (10% FBS) (Yang and Xiong 2012). In other words, the molar number of BSA in the supernatant is 5,000 times that of mAb. Considering that each antibody molecule has two identical antigen binding sites (IgM has 10 binding sites), and each BSA molecule has only one identical epitope. Therefore, BSA in the supernatant can fully saturate the binding sites of anti-BSA antibody. Considering the results of another experiment obtained (data not shown), there were many hybridomas secreting anti-OVA mAbs established by using SAL-OVA conjugate as immunogen. The results showed that using immunogen hapten-BSA conjugates as coating antigens can specifically screen out the hapten-specific hybridomas.

Due to the small size of hapten and the immunogenicity of cross linker, the bridging group connecting BSA and hapten may be involved in hybridoma screening. To avoid this situation, heterologous conjugates are always used as screening antigens in conventional hybridoma procedure (Esteve-Turrillas et al. 2010). However, the preparation of heterologous conjugates is usually laborious, time-consuming and expensive. More importantly, the coupling process may modify the antigenicity of haptens via influencing their original steric and electronic characteristics (Chappey et al. 1992), and thus reduce the screening efficiency of hapten-specific hybridoma. To avert such problems, an icELISA (as seen in this study) could be used to further confirm free hapten-specific hybridomas by using the hapten-BSA conjugate as a coating antigen and the hapten as a competitive inhibitor.

A blocking step is necessary to prevent any nonspecific binding of an antibody to the surface of microplates. If the blocking step can be omitted, the operation time and labor intensity of antigen-specific hybridoma screening can be further reduced. BSA is also a common blocking agent in ELISA methods (Lee et al. 2009). Therefore, when BSA is used as a carrier for immunogens, other blocking agents must be used, including casein, skimmed milk powder, and non-protein blocking solutions (such as Ficoll or polyvinyl alcohol) (Kim et al. 2004; López-Moreno et al. 2014; Huber et al. 2009). In this study, it was found that omitting the blocking step would not increase the nonspecific adsorption. The reason was presumably that the amount of BSA in the culture supernatant of hybridoma was far more than that of mAbs, which would give BSA more opportunities to occupy the residual active sites on the microwell surface.

## 5. Conclusion

The main disadvantages of the anti-hapten mAb preparation are the need to prepare heterologous conjugates and block microplates, which increase much workload and cost of the production, and even reduce the screening efficiency. As a carrier of haptens, this study showed the unique advantages of BSA in the development of haptic-specific mAbs, which greatly simplified the preparation process, reduced the preparation cost, and improved the screening efficiency of anti-hapten mAbs. In addition, the dcELISA is sensitive enough to be used for the determination of SAL in animal feed.

